# Neutrophil Extracellular Traps contribute to the pathogenesis of leprosy type 2 reactions

**DOI:** 10.1101/602722

**Authors:** Camila Oliveira da Silva, André Alves Dias, José Augusto da Costa Nery, Alice de Miranda Machado, Helen Ferreira, João Pedro Sousa Santos, Natalia Rocha Nadaes, Euzenir Nunes Sarno, Elvira Maria Saraiva, Verônica Schmitz, Maria Cristina Vidal Pessolani

## Abstract

Up to 50% of patients with the multibacillary form of leprosy is expected to develop acute systemic inflammatory episodes known as type 2 reactions (T2R), thus aggravating their clinical status. Thalidomide rapidly improves T2R symptoms. But, due to its restricted use worldwide, novel alternative therapies are urgently needed. A hallmark of T2R lesions is the presence of a neutrophil-rich inflammatory infiltrate. In this study, the potential involvement of neutrophil extracellular traps (NETs) production in T2R pathogenesis was investigated. Abundant NETs were found in T2R skin lesions, and increased spontaneous NETs formation was observed in T2R peripheral neutrophils. Both the *M. leprae* whole-cell sonicate and the CpG-Hlp complex, mimicking a mycobacterial TLR9 ligand, were able to induce NETs production *in vitro.* Moreover, TLR9 expression was shown to be higher in T2R neutrophils, suggesting that DNA recognition via TLR9 may be one of the pathways triggering this process during T2R. Finally, treatment of T2R patients with thalidomide for 7 consecutive days resulted in decreasing all of the evaluated *in vivo* and *ex vivo* NETosis parameters. Altogether, our findings shed light on the pathogenesis of T2R, which, it is hoped, will contribute to the emergence of novel alternative therapies in the near future.

**Author summary:** Leprosy is caused by a mycobacterium that has a predilection for skin and nerve cells. The chronic course of the disease may be interrupted by acute inflammatory episodes known as reactions, despite effective bacterial killing with antibiotics. Reactions aggravate patient’s clinical status and may become a medical emergency. Type 2 reactions (T2R) only occur in patients with high bacterial burden and is treated with thalidomide and/or corticosteroids. We are interested in understanding how inflammation is triggered and amplified during T2R. In this study we investigated the potential role of extracellular DNA released by neutrophils (known as NETs) in T2R, since they have been shown to cause inflammation. Abundant NETs were found in T2R skin lesions, and increased spontaneous NETs formation was observed in neutrophils present in the blood of T2R patients. Moreover, bacterial constituents were able to induce NETs production. Finally, treatment of T2R patients with thalidomide resulted in decreased NET formation. Altogether, our findings shed light on the pathogenesis of T2R, which, it is hoped, will contribute to the identification of biomarkers for early diagnosis and emergence of novel alternative therapies in the near future.

## Introduction

Leprosy, a disease widely associated with debilitating disfiguration, remains a public health threat in several low- and middle-income countries, including Brazil, with approximately 27,000 new cases each year [1]. The disease is caused by the obligate intracellular pathogen *Mycobacterium leprae* (ML) that targets skin macrophages and Schwann cells in the peripheral nerves as its preferential niche. Leprosy manifests as a spectrum of clinical forms in consonance with the immune response built by the host against the infection. In the multibacillary (MB) forms of the disease (lepromatous [LL], borderline lepromatous [BL]), patients generate weak specific cellular immunity, resulting in uncontrolled ML proliferation, numerous lesions, and extensive skin and nerve infiltration [2]. The chronic course of MB leprosy may be interrupted by acute inflammatory episodes known as type 2 reactions (T2R), the primary causes of peripheral nerve damage. Affecting 30-50% of these patients, erythema nodosum leprosum (ENL) presents as the most frequent manifestation of T2R. Characterized by painful cutaneous nodules or lesions, fever, joint and bone pain, iridocyclitis, neuritis, dactylitis, lymphadenitis, orchitis, and nephritis, and, in several aspects, often resembling typical chronic autoimmune diseases such as systemic lupus erythematosus (SLE) and rheumatoid arthritis, T2R presents as a systemic inflammation whose clinical symptoms range from mild to severe [3–5].

LL leprosy and a bacterial index (BI) of ≥ 4 are risk factors for developing T2R. While patients may present T2R as their first manifestation of leprosy, it is most frequently observed after the initiation of anti-mycobacterial multidrug therapy (MDT). T2R often runs a recurrent or chronic course, sometimes for many years, despite successful completion of MDT. In many countries, steroids are used to treat T2R. High doses are often required for long periods of time even though they do not always control the inflammation. In contrast, because treatment with thalidomide rapidly improves the major T2R symptoms, it is considered the drug of choice. But, due to its teratogenic effects, Brazil is the only country in which it is allowed [4,6].

Nonetheless, to this day, the T2R triggering mechanisms and immune-inflammatory pathways involved in its pathology remain ill defined. T2R is characterized by elevated levels of pro-inflammatory cytokines both at the lesion sites and systemically. The possible roles played by immune complex deposition and complement system activation in addition to that of specific cell-mediated immunity have been described [6,7]. Based on current evidence, it is reasonable to hypothesize that neutrophils play a central role in T2R pathogenesis. In contrast to non-reactional MB patients, T2R patients classically show an intense perivascular neutrophil infiltrate throughout the dermis and subcutis accompanied by neutrophilia [8]. Likewise, T2R peripheral neutrophils have been shown to display an activated phenotype in light of higher CD64 level expression [9] and the release of TNF [10] and pentraxin [11]. However, the precise mechanisms by which these cells contribute to the uncontrolled inflammation observed in T2R episodes remain nebulous.

In a recent report, we defined the recognition of nucleic acids by TLR9 as a major innate immunity pathway that is activated during T2R. Both ML and endogenous DNA-histone complexes were found at higher levels in the serum of T2R patients [12]. DNA recognition has been described as a major inflammatory pathway in several autoimmune diseases such as SLE whereas neutrophil DNA extracellular traps (NETs) have been shown to be a prime source of endogenous DNA [13]. Considering that neutrophil abundance is a marked characteristic of T2R lesions, the objective of this study was to investigate NETs production in T2R patients based on the hypothesis that the excessive NETs formation will trigger the activation and amplification of the immune-inflammatory pathways enrolled in its physio-pathogenesis.

## Methods

### Patient Recruitment

Leprosy patients were recruited at the Souza Araújo Outpatient Unit (Reference Center for Leprosy Diagnosis and Treatment, Laboratório de Hanseníase, Fundação Oswaldo Cruz, Rio de Janeiro, RJ, Brazil) from October 2016 thru December 2018. The patients were classified on the leprosy spectrum clinically and histologically based on Ridley–Jopling classification schemes [14] and were administered WHO-recommended leprosy multidrug treatment (MDT). The study population consisted of 2 groups of patients: i) LL/BL patients recruited before the start of MDT with no signs of reaction at the time of leprosy diagnosis (n=17: 11 LL and 6 BL, 10 men, 7 women, aged 22 – 80 with a median age of 50); and ii) T2R patients (n=23: 22 LL and 1 BL, 17 men, 6 women, aged 24 – 76 with a median age of 40) recruited at diagnosis of reaction. According to their clinical and histopathological findings, T2R manifested as erythema multiphorme (EM) in 4 and ENL in 19 of these patients [15]. Reaction occurred before (n=3), during (n=9), or after ending MDT (n=11). EM and ENL patients were treated with thalidomide at 100-300 mg/day and 16 were recruited for a reevaluation at day 7 of treatment (T2R_thal_). Healthy donors (HD) were also included (6 men, 10 women, aged 21 – 64, with a median age of 27) and participated in different assays during the study. None of the T2R patients had been treated with corticosteroid and/or thalidomide for at least 4 months prior to recruitment.

### Neutrophil isolation

Human neutrophils were isolated under endotoxin-free conditions from heparinized venous blood after density gradient centrifugation by Ficoll-Paque (GE Healthcare Life Sciences, NJ, USA). The neutrophil-rich layer was collected and residual erythrocytes were removed by lysis with an ACK (Ammonium-Chloride-Potassium) buffer. The quality of the neutrophil preparations was assessed by light microscopy of cytocentrifuged preparations, as previously described [10], and by flow cytometry using a mAb against CD16 conjugated with PE-Cy7 (catalog number 25-0168-42/clone CB16; BD Bioscience, CA, USA). The cells were assessed via the FACS Accuri flow cytometer (BD Bioscience); and the resulting data were analyzed by way of FlowJo V10 software (Tree Star). Neutrophil viability was estimated by way of Trypan blue-dye exclusion.

### Cell culture and neutrophil stimulation

Purified neutrophils (> 95% of the cells) were resuspended in RPMI medium 1640 (GIBCO^®^ Life technologies, MA, USA) and (1×10^6^) incubated at 37°C in centrifuge microtubes (Eppendorf, NY, USA) with or without a *M. leprae* whole-cell sonicate (MLWCS; 20 μg/ml; NR-19329; Bei Resources, VA, USA) or CpG-Hlp complex (rHlp: 0.25 μM; CpG 2395: 0.5 μM; Invivogen, CA, USA). The preparation of the CpG-Hlp complex was performed as previously described [12]. In the case of *in vitro* treatment with thalidomide (Abcam, CB, UK), the drug was dissolved in DMSO (Sigma-Aldrich, MO, EUA) and used at a final concentration of 50 μg/ml. NETs were quantified in the culture supernatant via the Quant-iT™ PicoGreen^®^ dsDNA Assay kit (Thermo Fisher Scientific, MA, USA) according to the manufacturer’s instructions. *In vitro* thalidomide efficacy was tested in human monocytes stimulated with LPS (Sigma-Aldrich), as previously described [16].

### Immunofluorescence

Immunofluorescence staining in frozen skin tissue was performed, as previously described [12]. For *in vitro* immunofluorescence staining, neutrophils (2×10^6^) were incubated in 24-well plates, adhered to poly-L-lysine-coated coverslips (1:10, Sigma-Aldrich), with or without MLWCS or CpG-Hlp, for 90 min and fixed with 4% paraformaldehyde. NETs were evidenced by DAPI (Sigma-Aldrich), rabbit polyclonal anti-MPO antibody (1:50 for *in vitro* immunofluorescence; 1:100 for tissue; catalog number sc-16128-R; Santa Cruz Biotechnology, CA, EUA), and mouse monoclonal anti-DNA/Histona H1 antibody (1:100; catalog number 05-457/clone AE-4; Merck Millipore, MA, EUA). Alternatively, the sections were then incubated with secondary antibodies: IgG anti-rabbit conjugated to Alexa Fluor 488^®^ and IgG2a anti-mouse conjugated to Alexa Fluor 633^®^ for immunofluorescence *in vitro* and IgG anti-rabbit conjugated to Alexa Fluor 633^®^ and IgG 2a anti-mouse conjugated to Alexa Fluor 594^®^ for tissue (Molecular Probes, OR, USA). Secondary antibodies were used at the concentration of 1: 1000. Coverslips were mounted with Permafluor (Thermo Scientific), sealed with mounting medium (Permount™ Mounting Medium; Fisher Chemical™/ Fisher Scientific), and analyzed via an AxiObserver Z1 Colibri microscope (Zeiss, NI, Germany). Images were processed by AxioVision software (Zeiss).

### TLR9 expression

Neutrophils (5 × 10^5^) were fixed with 4% paraformaldehyde and stored refrigerated (4°C) until use. The cells were centrifuged, suspended in PBS containing 1% BSA and an Fc receptor-blocking (human TruStainFcX; Biolegend, CA, USA). The neutrophils were then permeabilized with 0.1% saponin and incubated with a mAb against TLR9 conjugated with FITC (catalog number ab134369/clone 26C593.2; Abcam) or an isotype control and labeling assessed using FACS Accuri flow cytometer (BD Bioscience). The resulting data were analyzed by FlowJo V10 software (Tree Star).

### Quantification of DNA-histone and -MPO complexes

The levels of human histone (H1, H2A, H2B, H3, and H4)–associated DNA fragments in the serum were quantified by a photometric enzyme immunoassay (Roche Life Science, IN, USA); and analyzed as determined by the manufacturer.

For the quantification of the DNA-MPO complex levels in the serum samples, a previously described ELISA [17] was performed, with the exception of employing the anti-MPO antibody as capture, in which a different clone and supplier were used (catalog number sc-16128-R; Santa Cruz Biotechnology).

### Ethical considerations

This study was approved by the Ethics Committee of FIOCRUZ (CAAE 56113716.5.0000.5248). Informed written consent was obtained from all patients and healthy volunteers prior to specimen collection. All recruited individuals in this study were adults.

### Statistical analysis

Test results were represented as median. Statistically significant differences between the values were determined using the GraphPad Prism 5 (GraphPad Software Inc.), after the paired or unpaired t-test, or the parametric One-way analysis of variance (ANOVA) with a Bonferroni post-test. A p value < 0. 05 was considered significant.

## Results and discussion

The potential role of NETs in the pathogenesis of leprosy T2R was assessed in 23 T2R patients in comparison with 17 non-reactional LL/BL controls. Blood samples and biopsies of skin lesions were obtained. All patients recruited in this study were in attendance at the Outpatient Unit of the Oswaldo Cruz Foundation (Fiocruz), Rio de Janeiro, RJ, Brazil. LL/BL patients were recruited upon leprosy diagnosis and before initiating MDT. T2R patients were enrolled in the study at diagnosis of reaction and initiation of thalidomide treatment. Fifteen T2R patients were reassessed at day 7 of treatment (T2R_thal_). Healthy donors (HD) participated in a variety of assays during the study.

In most cases, the severe inflammatory process triggered during T2R leads to neutrophil infiltration [8]. Thus, as a first step, the presence of NETs in the skin lesions of T2R patients along with the potential effect of thalidomide on the abundance in NETs production were analyzed. Biopsies stained with H&E confirmed the classical presence of an important inflammatory neutrophil infiltrate in the T2R lesion, which, importantly, was drastically reduced by day 7 of treatment (T2R_Thal_) (S1 Fig). The lesions, labeled for NETs detection by immunofluorescence, utilized myeloperoxidase (MPO), histone H1, and DNA as markers. The co-localization of these molecules in filamentous structures characterizes NETs. It was observed that the intense NETs production detected in T2R patient skin lesions essentially disappeared as a result of thalidomide treatment (Fig 1A).

**Fig 1.**
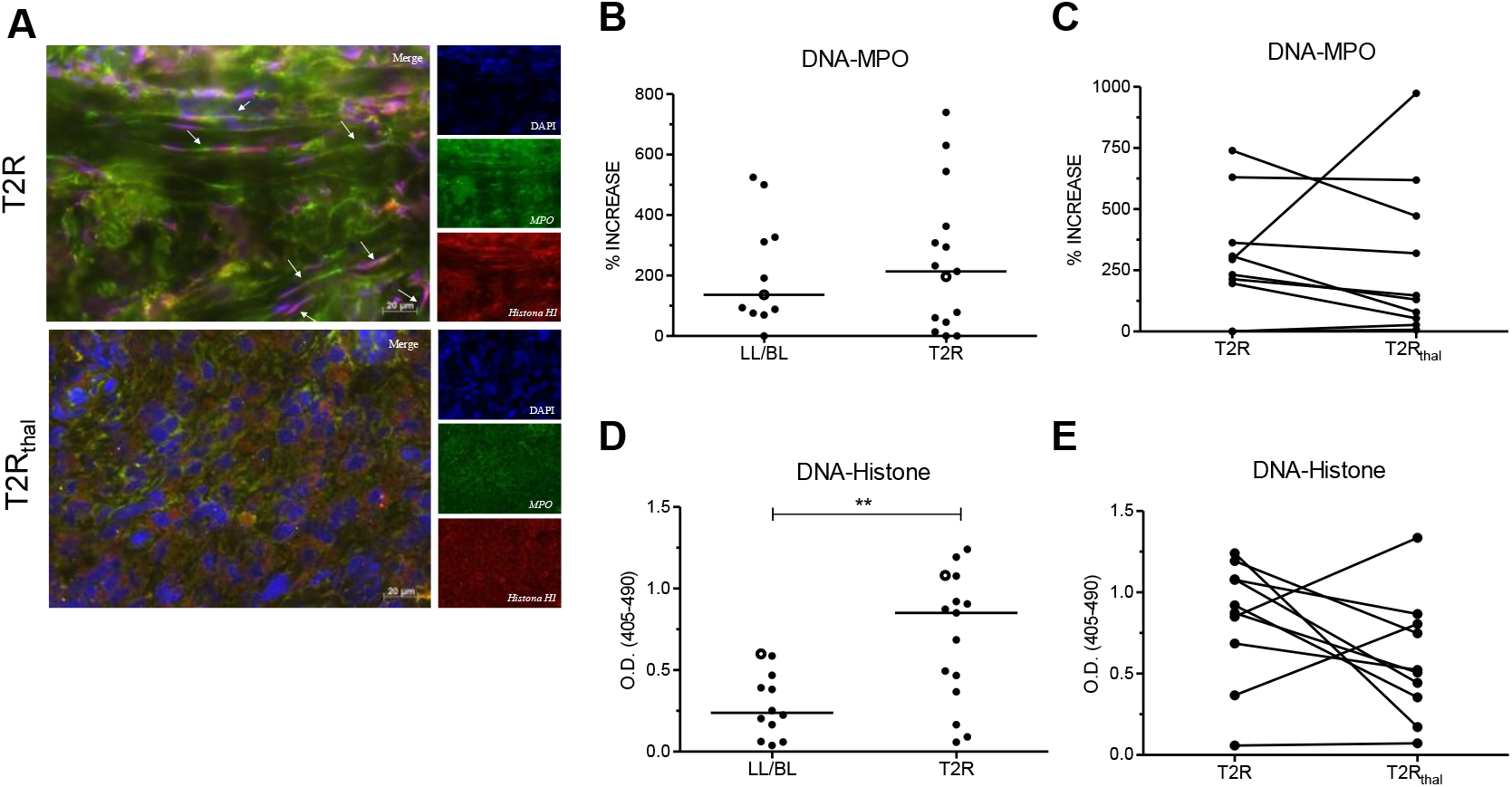
NETs are abundant in T2R skin lesions. (A) Skin lesions of T2R and T2R_thal_ patients were processed for immunostaining with antibodies against MPO (green) and histone (red), and DAPI was used for DNA staining (blue). The arrows indicate the presence of NETs characterized by a filamentous appearance and co-localization of these 3 macromolecules. The images are representative of 3 patients. Scale bar: 20 μm. (B – E) Patient serum samples were obtained; and the levels of the DNA-MPO and DNA-histone complexes were quantified by ELISA. Percentages of an increased DNA-MPO complex were obtained in relation to the blank. (C and E) Follow-up of the serum levels of the complexes during thalidomide treatment. Outliers were removed from analysis. Each dot represents a donor whereas the traits represent the median. The empty dots represent the patients shown in the immunofluorescence images. **p <0.01, T-test.

Circulating NETs-derived components have been demonstrated in several inflammatory diseases in which NETosis has been implicated in their pathogenesis [18–20]. The levels of MPO-or histone-DNA complexes in the serum of T2R patients were then investigated via specific immune enzymatic assays. A trend toward higher levels of the serum DNA-MPO complex was observed in T2R patients than those among non-reactional LL/BL (Fig 1B). Moreover, a longitudinal follow-up of 10 T2R patients showed a reduction in DNA-MPO complex levels in 7 of these patients at day 7 of treatment (T2R_thal_) (Fig 1C). Healthy donors presented lower levels of this complex, at a median value of 6.780% (IIQ: 0% – 54.64%; n=15), in relation to the blank. In confirmation of our previous data [12], significantly higher human DNA-histone complex levels were found in T2R than in LL/BL patients (Fig 1D). In addition, these same 7 out of the 10 T2R patients showed lower levels of this complex at day 7 of treatment (Fig 1E). The levels of this complex in healthy donors, however, were similar to the ones in the LL/BL group, at a median value of 0.3645 (IIQ: 0.1359 – 0.9568; n=15).

Peripheral neutrophils undergoing NETosis have been described in SLE, rheumatoid arthritis, and psoriasis [20–22]. So, as a next step, the production of NETs by T2R peripheral neutrophils in the absence of any stimulus was investigated. Blood was obtained, neutrophils were isolated, and their capacity to undergo *ex vivo* spontaneous NETosis was evaluated by DNA release in culture supernatants after a 90-min incubation period. Neutrophil preparations showed an average > 98% purity, as determined via microscopic analysis and flow cytometry (S2 Fig). When compared to neutrophils isolated from non-reactional LL/BL patients, higher levels of released DNA were detected in a T2R subgroup (Fig 2A). Interestingly, among the 7 patients with the highest DNA values, 5 developed reaction during MDT (red dots). Indeed, when this group was compared with the T2R patients who developed reaction before or after MDT, a significant difference was observed between them (Fig 2B). All T2R patients with the highest DNA levels showed reduced DNA release after 7 days of thalidomide medication (Fig 2C). Spontaneous *ex vivo* NETs formation was confirmed by fluorescence microscopy in neutrophils isolated from T2R patients as well as the reduction of NETs production upon administration of thalidomide (Fig 2D). This observation could explain the levels of NETs components in the serum of T2R patients, as shown in Fig 1. Altogether, these data indicate that excessive NETs formation occurs locally in the skin lesions and circulation of T2R patients and that this phenomenon tends to decline in thalidomide-treated patients.

**Fig 2.**
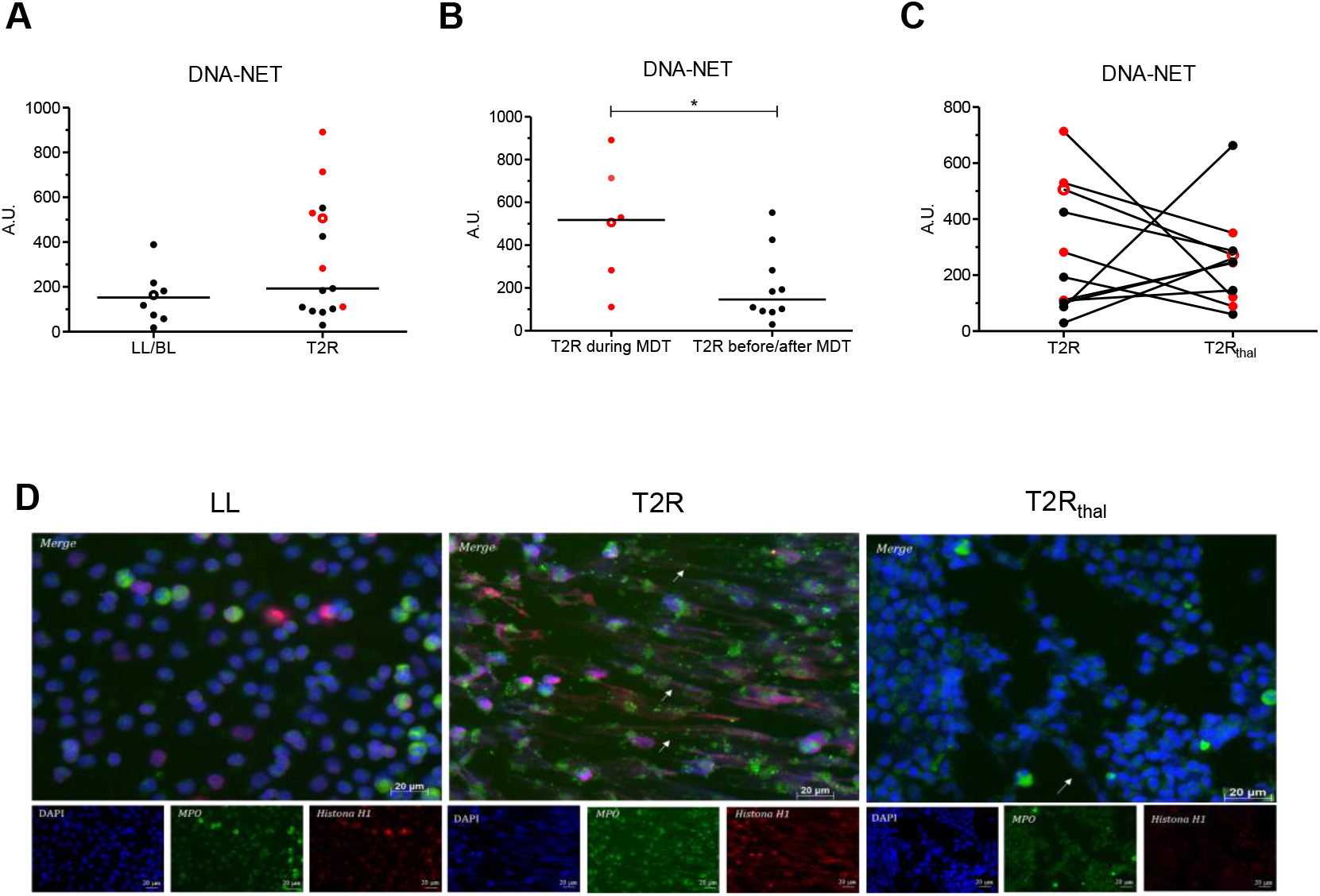
T2R neutrophils show spontaneous *ex vivo* NETs production, which decreases with thalidomide treatment. (A – C) Neutrophils from LL/BL, T2R, and T2R_thal_ patients were incubated for 90 min and DNA release was measured by picogreen staining. Each dot represents a donor. Red dots represent patients who developed T2R during MDT. The traits represent the median. The empty dots represent the patients shown in the immunofluorescence images. (B) Comparison of patients who developed T2R during MDT versus those with reactional episodes before or after MDT. (C) Follow-up of DNA levels released after 7 days of thalidomide treatment. (D) Immunofluorescence images show NETs labeled with antibodies against MPO (green) and histone (red), and DAPI staining DNA (blue). Images are representative of 6 LL/BL, 8 T2R and 8 T2R_thal_. Scale bar: 20 μm. *p <0.05, T-test.

The higher levels of spontaneous NETs formation observed in reactional MDT-T2R patients may be related to the massive release of ML components from infected tissue taking place during treatment. Since bacterial components are known to trigger NETosis [23], the capacity of the ML whole-cell sonicate (MLWCS) to induce NETs formation *in vitro* was investigated. S3A Fig outlines the dose-response curve of DNA release by neutrophils isolated from healthy donors stimulated for 90 min with MLWCS. The concentration of 20 μg/mL induced significantly higher levels of NETs production relative to those in non-stimulated cells and, as a result, was used in the remaining assays of the study. Time-course assays were also performed, indicating 90 min as the best time point for NETs formation in response to MLWCS (S3B Fig). S3C Fig shows the immunofluorescence image of MLWCS-induced NETs production, confirming the capacity of MLWCS to induce NETs formation in healthy donor neutrophils *in vitro.*

Consequently, the ability of MLWCS to induce NETs production in leprosy patient peripheral neutrophils was then examined. MLWCS stimulated the neutrophils of all 3 groups of patients to release DNA at higher levels than in the non-stimulated cultures (Fig 3A). NETs formation in response to MLWCS in neutrophils of leprosy patients was confirmed by immunofluorescence (Fig 3B).

**Fig 3.**
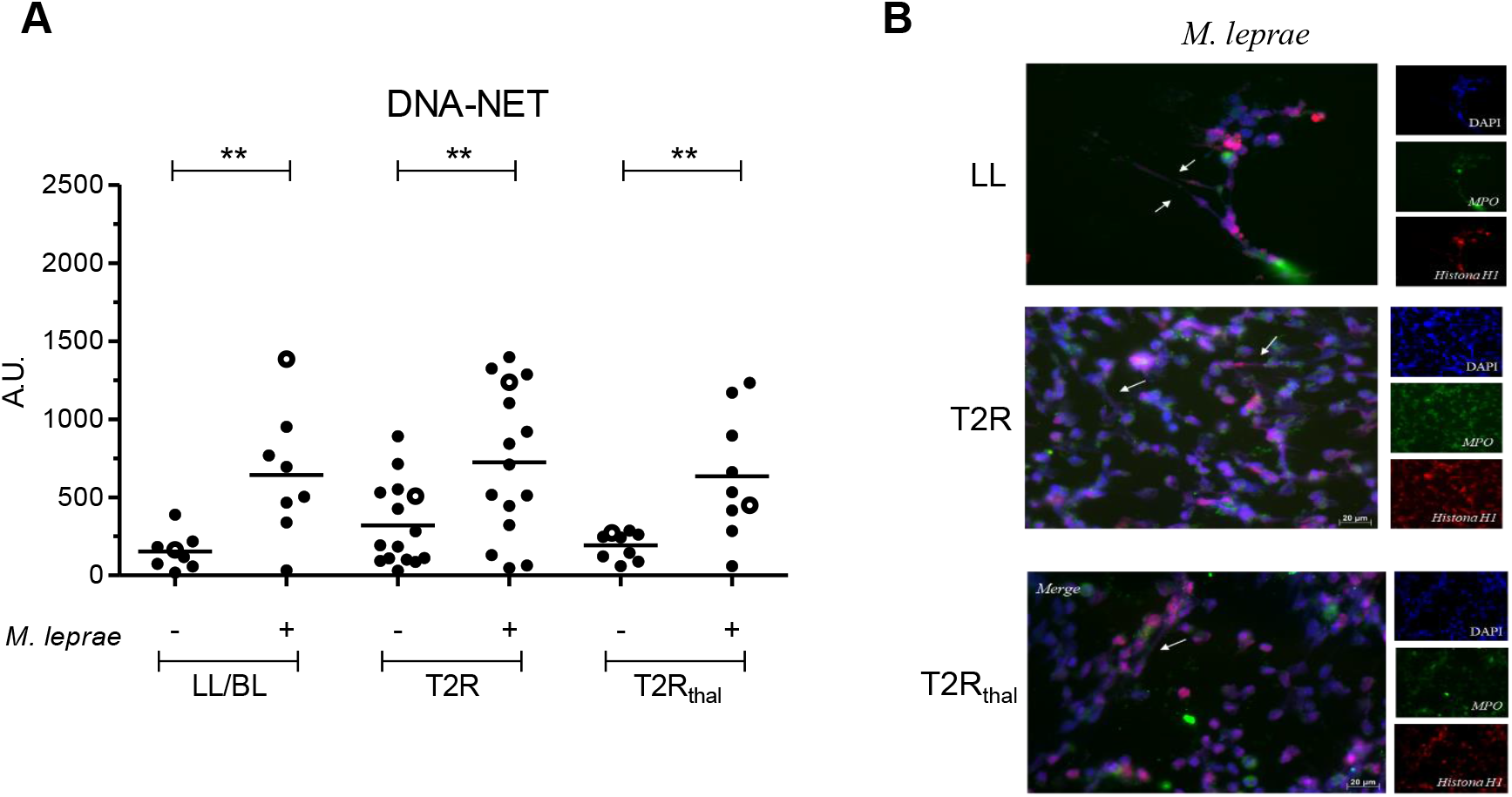
*M. leprae* induces NETs formation *in vitro.* Neutrophils from LL/BL, T2R, and T2R_thal_ patients were isolated and stimulated or not with *M. leprae* whole-cell sonicate (MLWCS). (A) DNA release was measured in culture supernatants by pycogreen staining. Each dot represents a donor; and the traits represent the median. The empty dots are the patients shown in the immunofluorescence images. (B) Immunofluorescence images show NETs labeled with antibodies against MPO (green), histone (red), and DAPI staining DNA (blue). Images are representative of 6 LL/BL, 8 T2R, and 8 T2R_thal_. The arrows indicate the presence of NETs, characterized by a filamentous appearance and co-localization of these 3 macromolecules. Scale bar: 20 μm. ** p < 0.01, T-test.

The apparently *in vivo* inhibitory effect of thalidomide on NETs production in T2R patients observed both at the lesion site (Fig 1A) and in peripheral neutrophils (Fig 2) raised the hypothesis that the drug might be acting directly in this process. As a test, healthy-donor neutrophils were stimulated *in vitro* with MLWCS in the presence or not of thalidomide while NETs production was monitored by DNA release. S4A Fig shows the results obtained with one donor, in which no effect of thalidomide on MLWCS-induced NET formation was observed. S4B Fig summarizes the results of 6 healthy donors, only one of whom experienced an inhibitory effect. As expected, however, thalidomide was able to block TNF secretion by monocytes in response to LPS, as previously referred to [16] (S4C Fig). The incapacity of thalidomide to directly interfere in the NETs production process is reinforced by the observation that T2R_thal_ neutrophils continued to respond to MLCWS *in vitro* even after receiving *in vivo* thalidomide (Figs 3A and 4A). These data suggest that the inhibitory *in vivo* effect of thalidomide on NETs formation is, albeit indirectly, most probably related to the capacity of this drug to block pro-inflammatory cytokine production such as IFN-I [24], a mediator known to prime neutrophils to NETosis [25].

**Fig 4.**
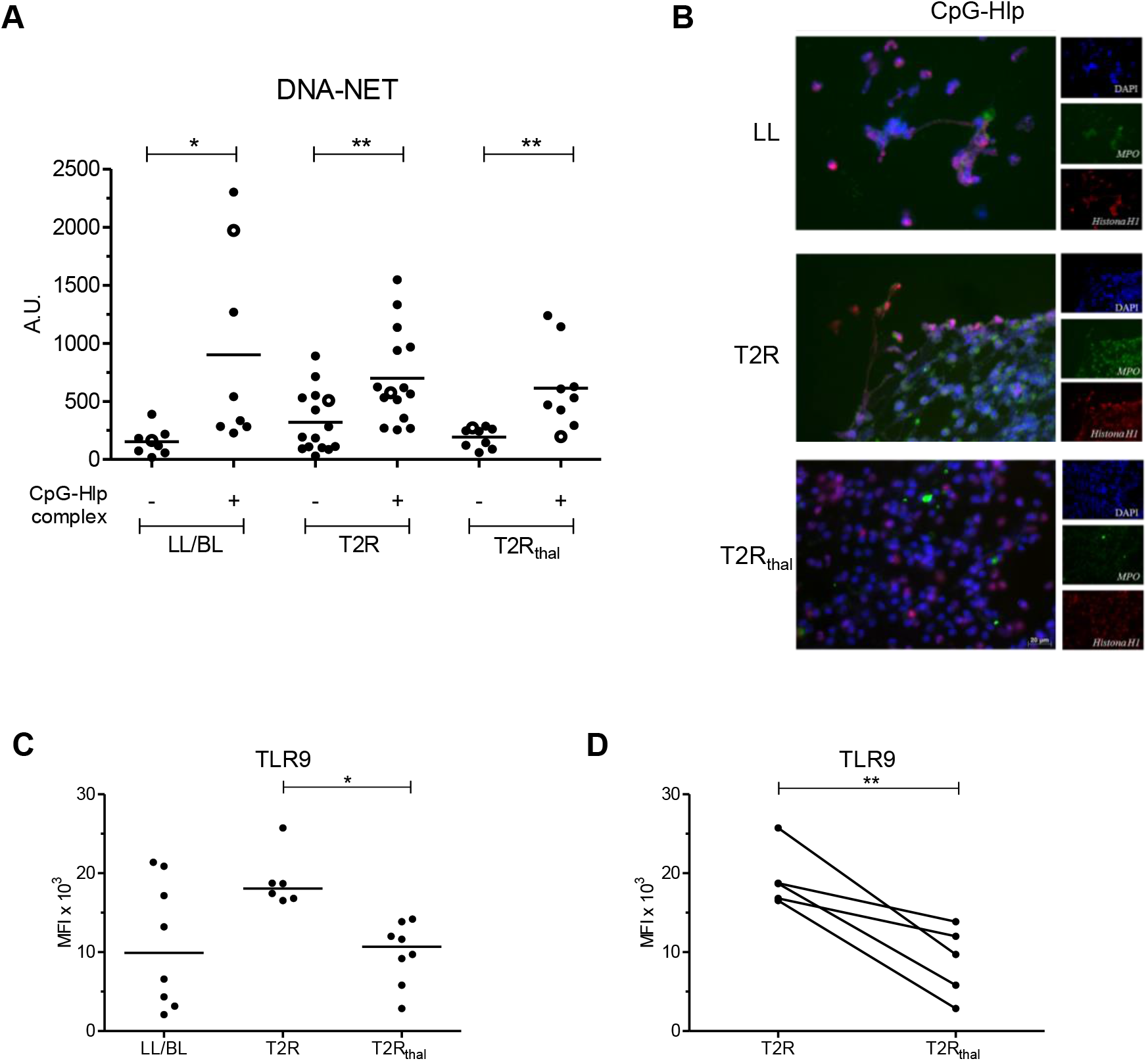
TLR9 ligands induce NETs formation; and T2R patient neutrophils express higher levels of this receptor. (A) Neutrophils isolated from LL/BL, T2R, and T2R_thal_ patients were stimulated or not with the CpG-Hlp complex; and DNA release was measured in culture supernatants by pycogreen. Each dot represents a donor, and the traits represent the median. The empty dots are the patients shown in the immunofluorescence images. (B) NETs were visualized by immunofluorescence after staining for MPO (green), histone (red), and DNA (blue). Scale bar: 20 μm. (C) Neutrophils of LL/BL, T2R and T2R_thal_ patients were isolated and *ex vivo* TLR9 expression levels were quantified by flow cytometry. (D) Follow-up of TLR9 expression in T2R patients after 7 days of thalidomide treatment. *p <0.05; ** p <0.01, T-test and one-way ANOVA with Bonferroni posttest correction.

TLRs had been previously implicated in NETosis induction [23]. In our previous study, besides the higher levels of endogenous histone-DNA complexes, higher levels of the mycobacterial histone-like protein (Hlp) in T2R patients sera, probably released as a consequence of bacterial killing during MDT [12], was also detected. Thus, we hypothesized that Hlp complexed to DNA, a TLR9 ligand, could be one of the bacterial components present in MLWCS that is responsible for NETs induction. To test this hypothesis, recombinant Hlp was obtained and then combined with the CpG oligonucleotide, mimicking bacterial DNA. Neutrophils were then stimulated with the CpG-Hlp complex for 90 min; and NETs production was monitored by measuring DNA release in culture supernatants. CpG-Hlp was shown to induce NET formation *in vitro* in leprosy patient neutrophils at levels comparable to those seen for MLWCS (Fig 4A). The capacity of CpG-Hlp to stimulate NETs production in leprosy patients was confirmed by immunofluorescence (Fig 4B).

We have previously demonstrated higher expression levels of TLR9 in skin lesions and T2R blood mononuclear cells as opposed to LL/BL patient levels [12]. Thus, as a next step, the status of TLR9 expression in leprosy patient neutrophils was compared. The median fluorescence intensity (MFI) values of TLR9 in T2R neutrophils were higher overall relative to those of LL/BL patients.

Moreover, a significant decrease in TLR9 expression was observed in T2R_thal_, reaching values close to the ones seem in T2R (Fig 4C). Indeed, during follow-up of 5 T2R patients, an accentuated drop in TLR9 expression after 7 days of thalidomide treatment was shown (Fig 4D). Representative histograms are exhibited in S5 Fig. It is noteworthy that although T2R and T2R_thal_ neutrophils express different TLR9 levels, both demonstrated a similar capacity to respond *in vitro* to CpG-Hlp. One possible explanation for this observation may be the high concentration of CpG-Hlp used in these *in vitro* assays, constituting a supra-optimal stimulation. The use of sub-optimal CpG-Hlp concentrations may disclose differences among the groups.

The capacity of CpG-Hlp to induce NETs formation *in vitro* together with the higher levels of TLR9 expression and the simultaneous increase in the circulating levels of its ligands (both the human DNA-histone and -Hlp complexes [12]) strongly suggest that DNA recognition via TLR9 is involved in NETs production during T2R. These parameters decreased after 7 days of treatment with thalidomide (Figs 1 and 4), which may explain the diminished spontaneous NETs formation observed in T2R_thal_ *in vivo* (Fig 2). It is worth mentioning again that thalidomide is an extremely effective drug in the remission of T2R clinical symptoms [26,27]. Thus, the blockage of NETs formation observed in treated patients reinforces the idea that excessive NETs generation may play an important role in T2R pathogenesis.

NETs have been implicated in the immunopathogenesis of SLE and other chronic inflammatory autoimmune diseases via the following mechanisms: 1) the externalization of modified autoantigens; 2) induction of type I IFN; 3) autoantibody production; 4) stimulation of the inflammasome; 5) activation of the classical and alternative pathways of the complement system; and 6) direct effects on the endothelium and the consequent induction of vasculopathy [28]. Since most of these events have also been shown to be activated during T2R [7], it is reasonable to speculate that the excessive NETs production herein described performs a central role in triggering and amplifying these pathological pathways in T2R. Novel therapies under development that target the accumulation of NETs [29] may prove useful in treating both autoimmune inflammatory diseases and T2R.

## Acknowledgments

We thank the Rede de Plataformas Tecnológicas Fiocruz (Rpt08a-RJ) for its support in the flow cytometry experiments and analyses, and Judy Grevan for editing the text.

## Supporting information Captions

**S1 Fig.**
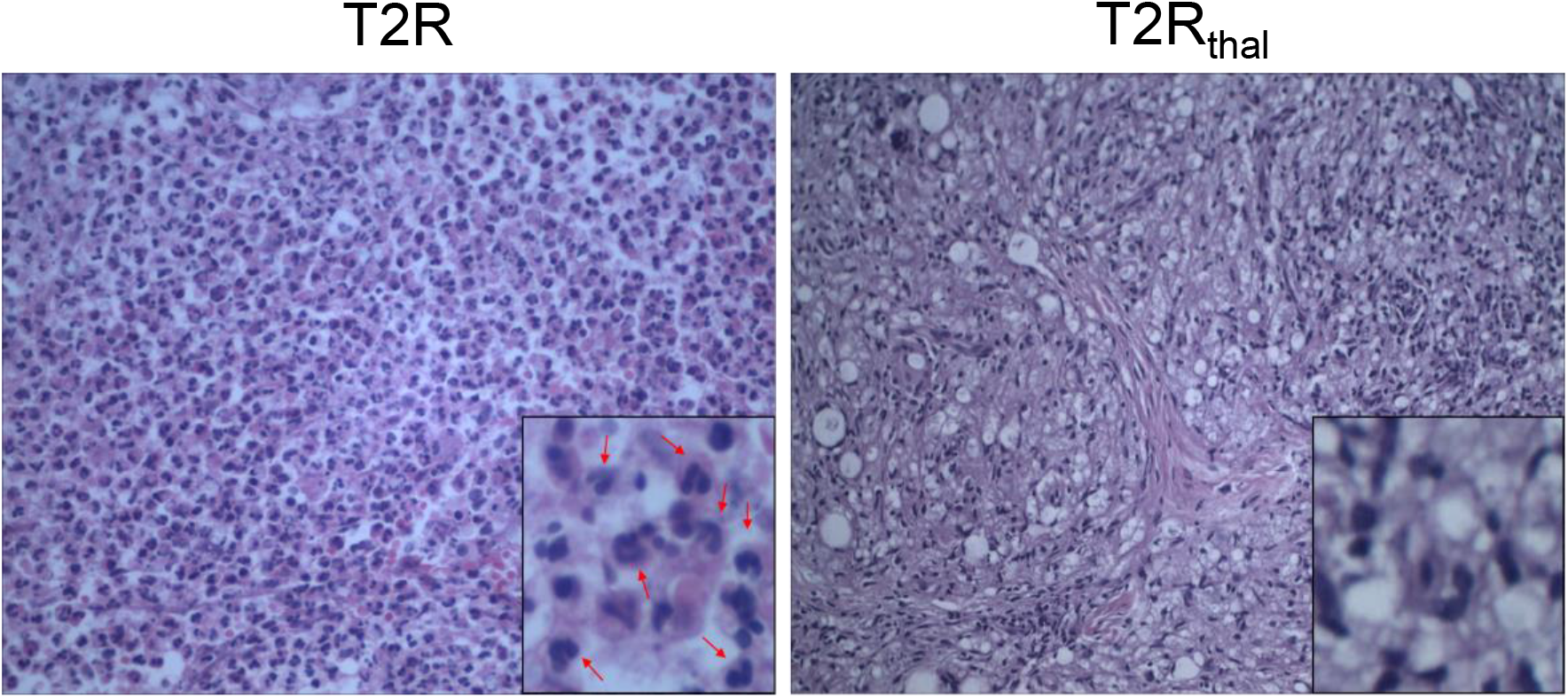
Neutrophils in T2R skin lesions. Histology of cutaneous lesions from T2R and T2R_thal_ patients stained by H&E. The arrows indicate the presence of neutrophils. Representative images of 3 patients (400x).

**S2 Fig.**
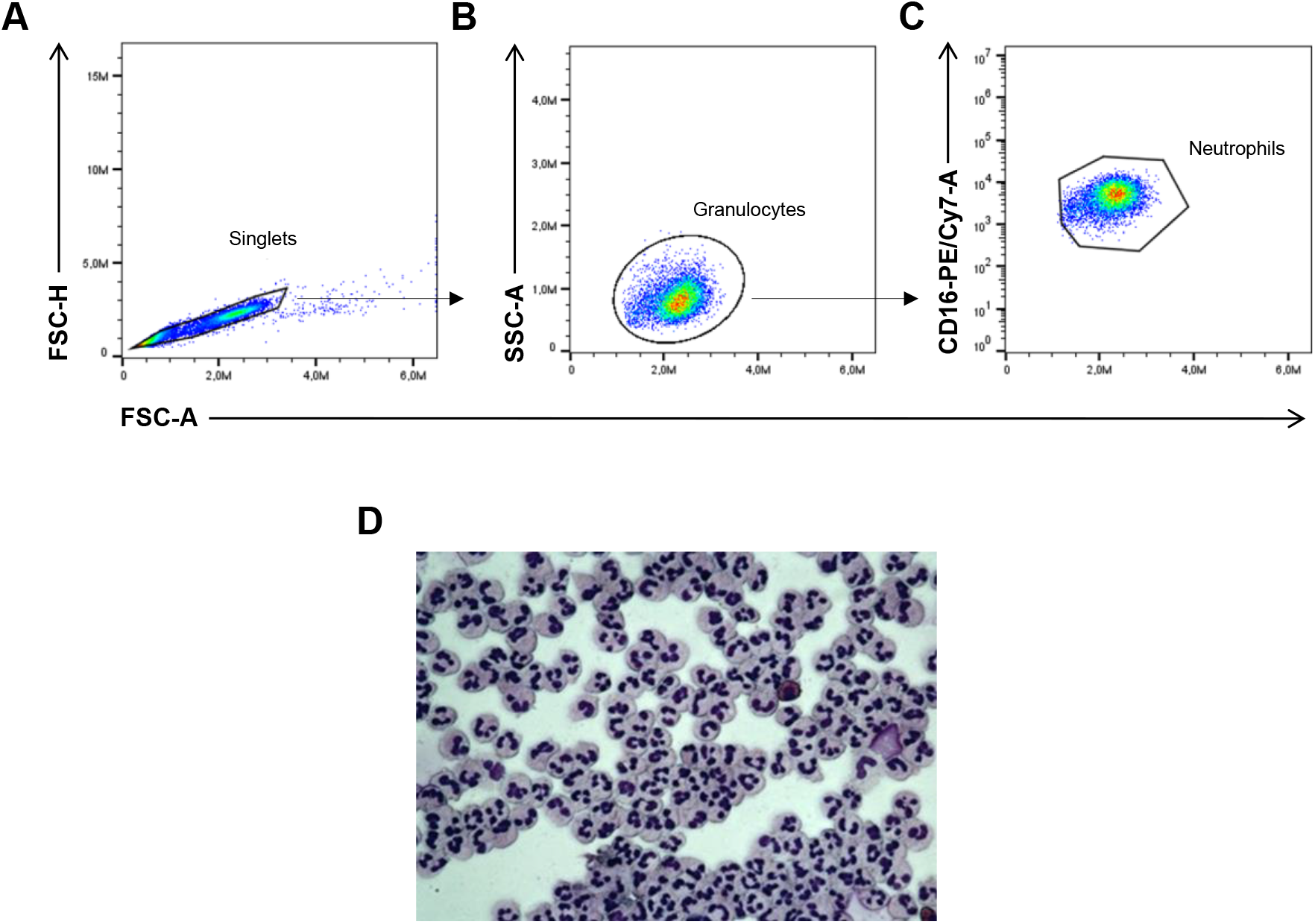
Rate of purity of isolated neutrophils. (A – C) Representative dot-plot diagram of the flow cytometry of neutrophils isolated from healthy donors for rate-of-purity evaluation. (A) *Gate* generated by FSC-A vs. FSC-H parameters for singlet analysis. (B) The granulocytic population *gate* was generated by FSC-A vs. SSC-A parameters. (C) The percentage of CD16+ cells was provided by gating FSC-A *vs.* CD16 axes. Representative of 4 healthy donors. (D) Representative image of a cytospin slide of purified healthy donor neutrophils (n = 4).

**S3 Fig.**
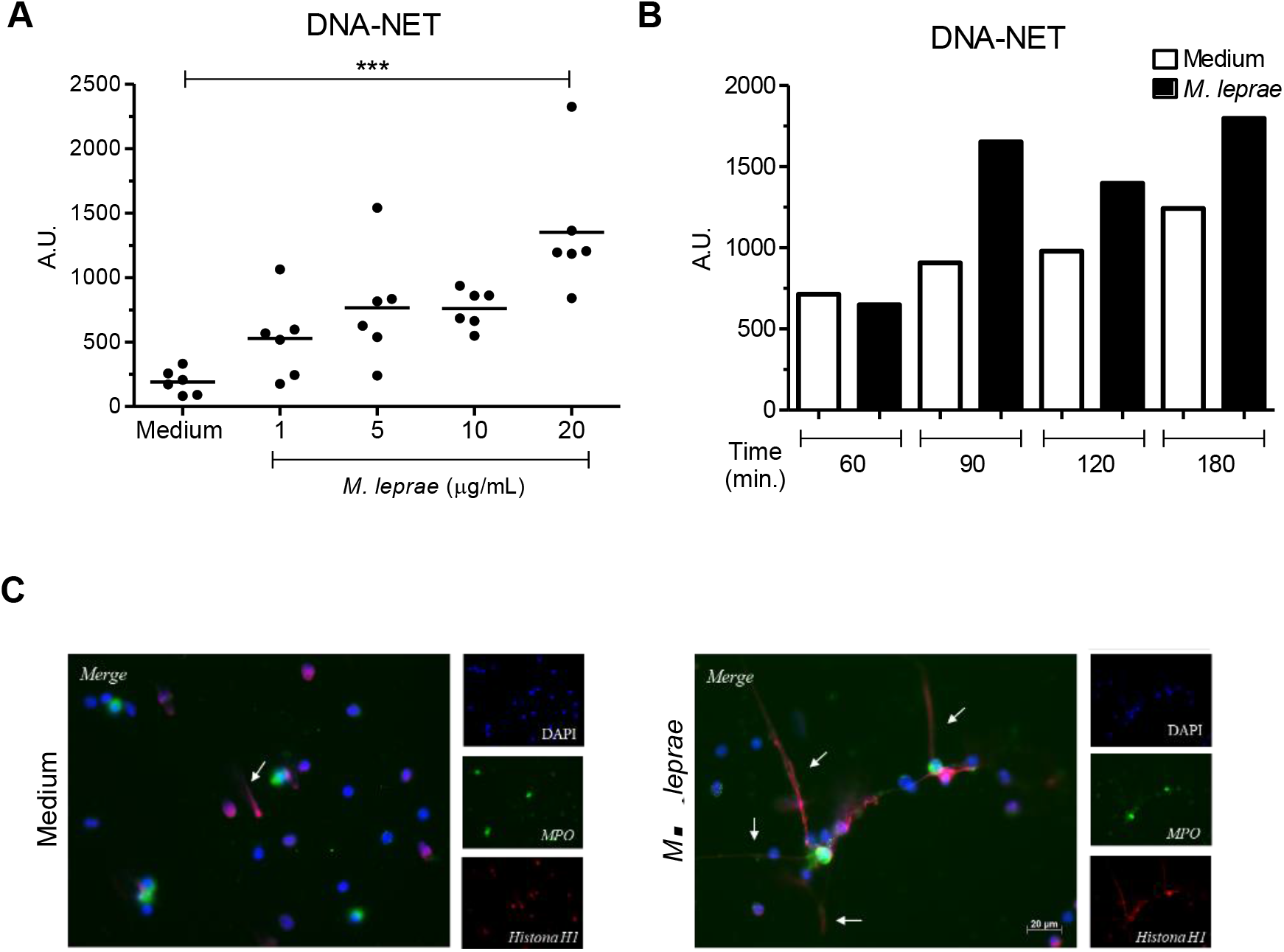
*M. leprae* induces NET formation. (A) Healthy donor neutrophils (1 × 10^6^) were stimulated or not with 20 μg/mL of a *M. leprae* whole-cell sonicate (MLWCS) for 60, 90, 120, and 180 min; and DNA release was measured by picogreen. Representative of 3 healthy donors. (B) Neutrophils (1×10^6^) from healthy donors (n=6) stimulated or not with MLWCS at different concentrations (1, 5, 10, and 20 μg/mL) for 90-min incubation. DNA release in the supernatant was measured by picogreen. Each dot represents a donor and the traits represent the median. (C) Neutrophils (2×10^6^ cells) from healthy donors were stimulated or not with 20 μg/mL of MLWCS for 90 min. Representative immunofluorescence image of 4 healthy donors. ***p<0.001, one way ANOVA with Bonferroni post-test.

**S4 Fig.**
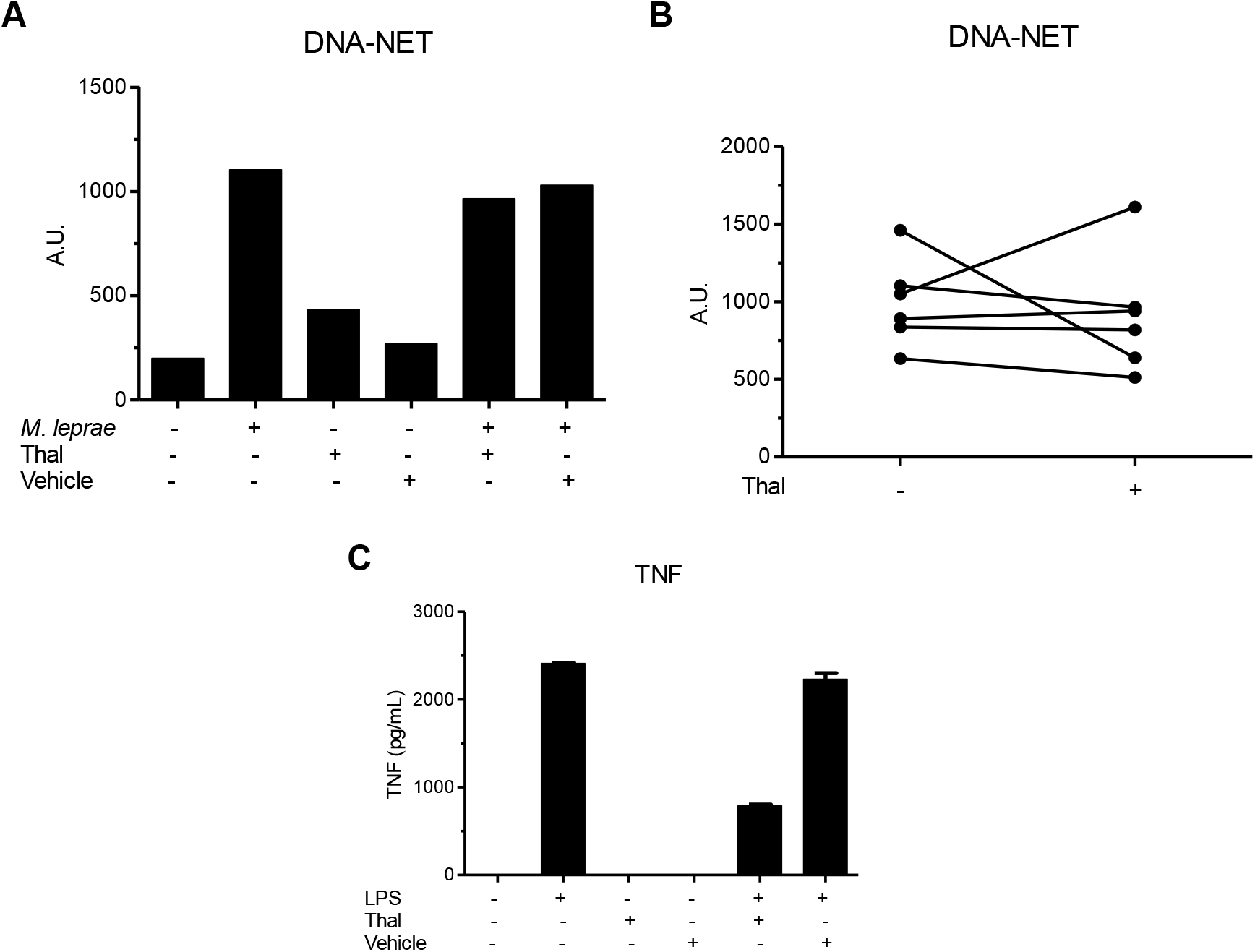
Fig. *In vitro* effect of thalidomide on *M. leprae-induced* NETs formation. (A) Neutrophils (1×10^6^ cells) from healthy donors were stimulated or not with MLWCS (20 μg/mL) and/or thalidomide (50 μg/mL) for 90-min incubation; and DNA release was measured by picogreen. Representative of 6 healthy donors. (B) Follow-up of DNA release by healthy donor neutrophils (n =6) stimulated with MLWCS in the presence or absence of thalidomide. Each dot represents a donor. (C) To test the efficacy of *in vitro* thalidomide, monocytes (2×10^6^ cells) from healthy donors were stimulated or not with LPS (1 μg/mL) and/or thalidomide (50 μg/mL) for an 18h-incubation period for TNF release dosing by ELISA. Data represent median of 2 healthy donors. DMSO was used as vehicle.

**S5 Fig.**
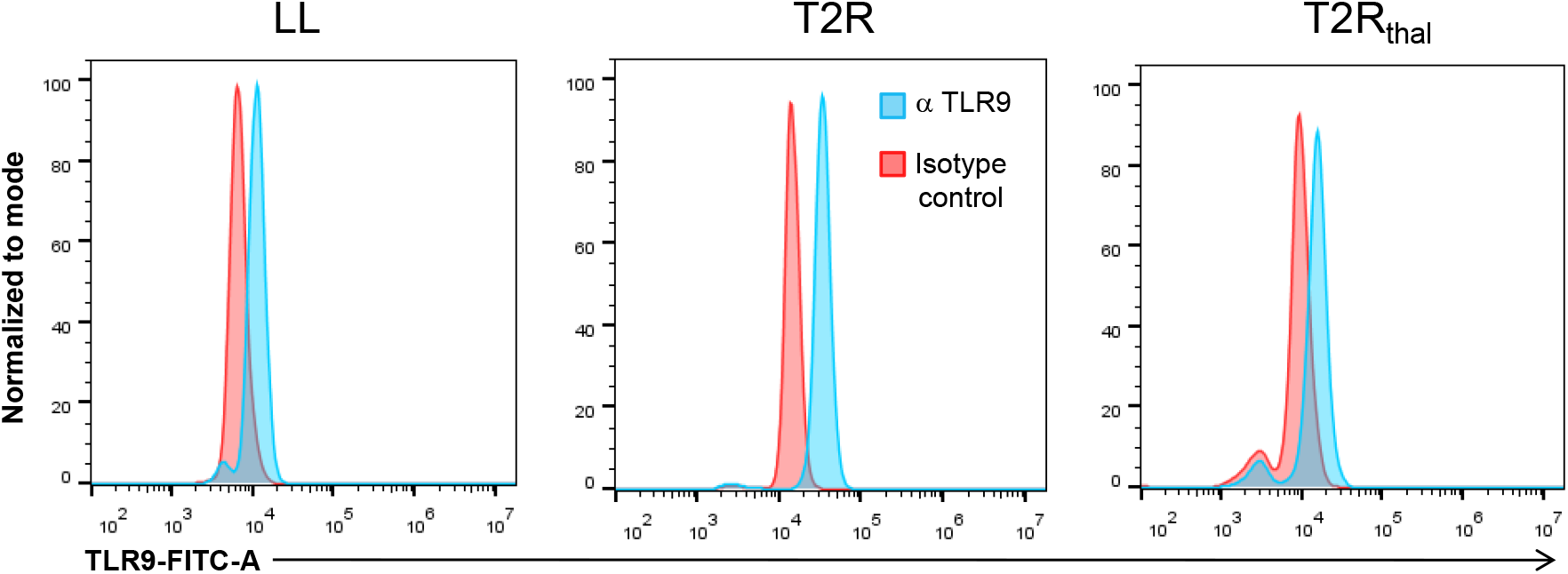
Levels of TLR9 expression in leprosy patient neutrophils. Representative histograms showing the quality of anti-TLR9 antibody labeling in neutrophils isolated from the different groups of analyzed patients.

